# Immune Evasion and Membrane Fusion of SARS-CoV-2 XBB Subvariants EG.5.1 and XBB.2.3

**DOI:** 10.1101/2023.08.30.555188

**Authors:** Julia N. Faraone, Panke Qu, Negin Goodarzi, Yi-Min Zheng, Claire Carlin, Linda J. Saif, Eugene M. Oltz, Kai Xu, Daniel Jones, Richard J. Gumina, Shan-Lu Liu

**Affiliations:** Center for Retrovirus Research, The Ohio State University, Columbus, OH 43210, USA; Department of Veterinary Biosciences, The Ohio State University, Columbus, OH 43210, USA; Molecular, Cellular, and Developmental Biology Program, The Ohio State University, Columbus, OH 43210, USA; Department of Internal Medicine, Division of Cardiovascular Medicine, The Ohio State University, Columbus, OH 43210, USA; Center for Food Animal Health, Animal Sciences Department, OARDC, College of Food, Agricultural and Environmental Sciences, The Ohio State University, Wooster, OH 44691, USA; Veterinary Preventive Medicine Department, College of Veterinary Medicine, The Ohio State University, Wooster, OH 44691, USA; Viruses and Emerging Pathogens Program, Infectious Diseases Institute, The Ohio State University, Columbus, OH 43210, USA; Department of Microbial Infection and Immunity, The Ohio State University, Columbus, OH 43210, USA; Department of Pathology, The Ohio State University Wexner Medical Center, Columbus, OH, USA; Dorothy M. Davis Heart and Lung Research Institute, The Ohio State University Wexner Medical Center, Columbus, OH 43210, USA; Department of Physiology and Cell Biology, College of Medicine, The Ohio State University Wexner Medical Center, Columbus, OH 43210, USA

## Abstract

Immune evasion by SARS-CoV-2 paired with immune imprinting from monovalent mRNA vaccines has resulted in attenuated neutralizing antibody responses against Omicron subvariants. In this study, we characterized two new XBB variants rising in circulation — EG.5.1 and XBB.2.3, for their ability of neutralization and syncytia formation. We determined the neutralizing antibody in sera of individuals that received a bivalent mRNA vaccine booster, BA.4/5-wave infection, or XBB.1.5-wave infection. Bivalent vaccination-induced antibodies neutralized efficiently ancestral D614G, but to a much less extent, two new EG.5.1 and XBB.2.3 variants. In fact, the enhanced neutralization escape of EG.5.1 appeared to be driven by its key defining mutation XBB.1.5-F456L. Notably, infection by BA.4/5 or XBB.1.5 afforded little, if any, neutralization against EG.5.1, XBB.2.3 and previous XBB variants — especially in unvaccinated individuals, with average neutralizing antibody titers near the limit of detection. Additionally, we investigated the infectivity, fusion activity, and processing of variant spikes for EG.5.1 and XBB.2.3 in HEK293T-ACE2 and CaLu-3 cells but found no significant differences compared to earlier XBB variants. Overall, our findings highlight the continued immune evasion of new Omicron subvariants and, more importantly, the need to reformulate mRNA vaccines to include XBB spikes for better protection.

## Introduction

The COVID-19 pandemic still lingers across the globe as its causative agent, severe acute respiratory syndrome virus 2 (SARS-CoV-2), continues to evolve. This evolution challenges the efficacy of current vaccines, requiring the constant surveillance and reassessment of current public health measures against COVID-19. Since the emergence of the Omicron lineage of SARS-CoV-2 in 2022, the virus has exhibited ever-increasing numbers of mutations that escape neutralizing antibodies generated through both mRNA vaccination and SARS-CoV-2 convalescence^1-8^. The XBB-lineage subvariants, which evolved from the recombinant XBB variant in early 2023, have displayed particularly strong immune escape^3,5,7,9-18^. This new level of immune evasion has prompted the Food and Drug Administration to recommend inclusion of XBB-lineage subvariants in future iterations of mRNA vaccines^19^.

One concern in vaccine design is the role of immune imprinting, which impairs vaccine efficacy against evolving variants. It has been demonstrated that the three-dose course of wildtype spike mRNA vaccine may be biasing immune responses toward earlier lineages of the virus, impairing our ability to mount effective responses toward more recent Omicron-lineage subvariants^20-22^. The bivalent booster dose, including both the wildtype and BA.4/5 spikes, augments the response toward Omicron subvariants relative to the 3-dose course of monovalent vaccines, but only to a limited extent^7,20,21^. Additional doses of Omicron spike-based vaccines or exposure to Omicron-lineage variants has been shown to more effectively counteract immune imprinting, suggesting the need to reconfigure current approaches^20^. The continued surveillance and characterization of emerging variants is critical for informing such decisions.

This study focuses on two XBB-lineage variants currently on the rise, termed EG.5.1 and XBB.2.3^23,24^. The latter evolved directly from XBB, with two additional mutations in spike: D253G in the N-terminal domain (NTD) and P521S in the receptor binding domain (RBD). EG.5.1 evolved from XBB.1.5, with two additional mutations in spike: Q52H in the NTD and F456L in the RBD^25^ (**Fig 1A**). EG.5.1, in particular, has increased rapidly in circulation across the globe and is currently on track to become a dominant variant^24^. Our study sought to characterize these variants and their defining mutations by investigating aspects of spike protein biology, including infectivity, fusogenicity, and escape from neutralizing antibodies in bivalent vaccinated sera, BA.4/5-wave convalescent sera, and XBB.1.5-wave convalescent sera, as well as the monoclonal antibody (mAb) S309. We compare these attributes to spikes from the ancestral D614G and late-evolved Omicron subvariants BA.4/5, XBB, XBB.1.5, and XBB.1.16.

**Figure 1:**
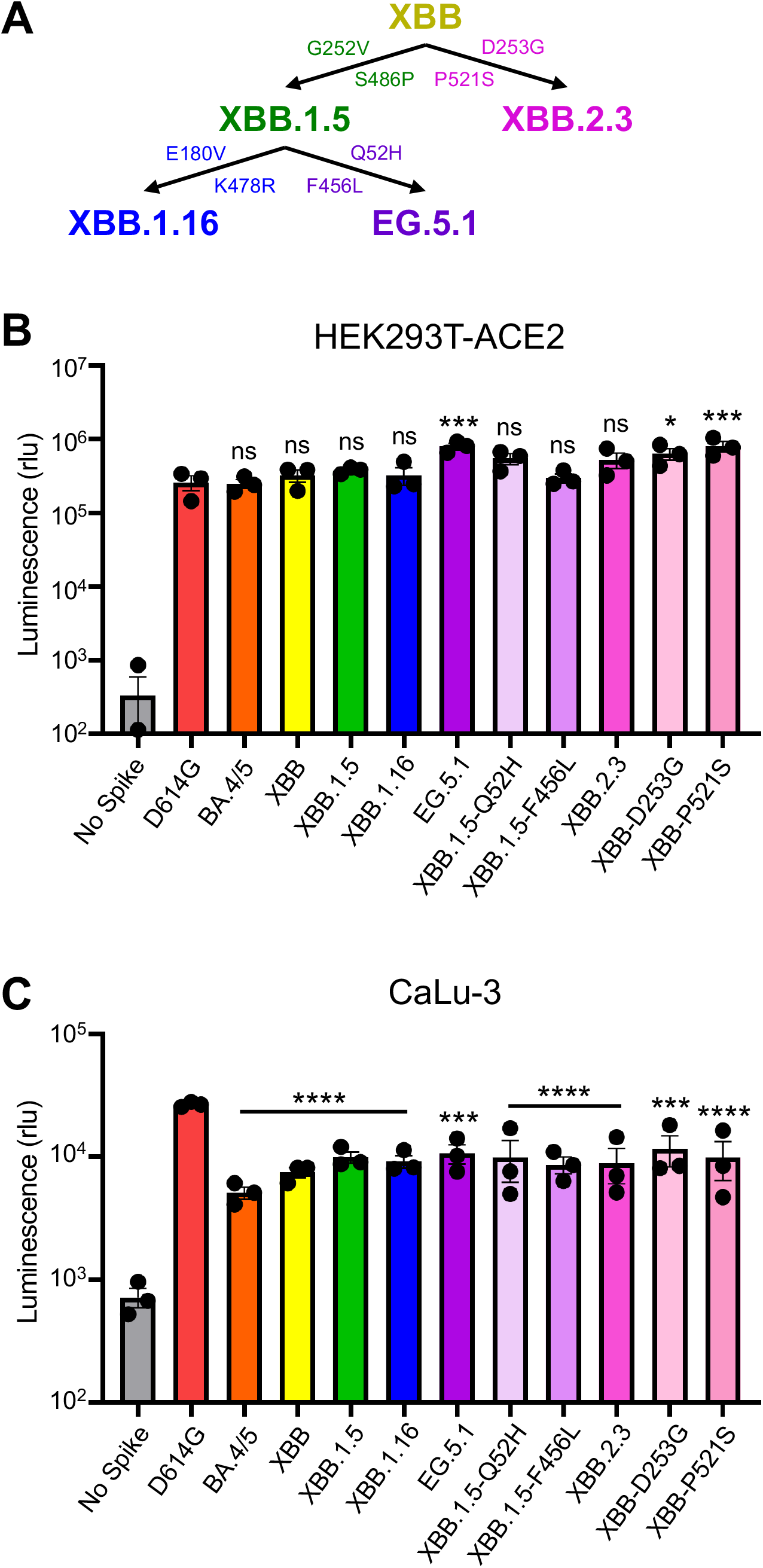
Infectivity of pseudotyped lentiviruses bearing XBB.2.3 or EG.5.1 spike into HEK293T-ACE2 and CaLu-3 cells. **(A)** Schematic relationship between XBB-lineage variants in this study. Arrows denote direct relationships between variants with the corresponding spike mutations written along them. **(B and C)** Pseudotyped lentiviruses bearing each of the depicted spikes of interest were produced in HEK293T cells and used to infect **(B)** HEK293T-ACE2 or **(C)** CaLu-3 cells. Bars in **(B and C)** represent means ± standard deviation for 3 replicates represented by individual dots (n=3). p values are displayed as *p < 0.05, ***p < 0.001, ****p < 0.0001 and ns p > 0.05.

## Results

### EG.5.1 and XBB.2.3 have comparable infectivity in HEK293T-ACE2 and CaLu-3 cells

We first determined the infectivity of pseudotyped lentiviruses bearing each spike in HEK293T cells expressing human ACE2 (HEK293T-ACE2), as well as the human lung epithelial carcinoma cell line CaLu-3. In HEK293T-ACE2 cells, EG.5.1 exhibited slightly higher infectivity relative to the parental subvariant XBB.1.5, with a 2.1-fold increase (p < 0.05) (**Fig 1B**). This enhancement appears to be largely driven by the XBB.1.5-Q52H single mutation, which exhibited a 1.4-fold increase relative to XBB.1.5 (p > 0.05), while XBB.1.5-F456L alone did not cause any increase in infectivity relative to XBB.1.5 (**Fig 1B**). While XBB.2.3 exhibited comparable infectivity relative to XBB (p > 0.05), two single mutations, XBB-D253G and XBB-P521S, conferred an increased titer relative to XBB of 2.5-fold (p > 0.05) and 1.7-fold (p < 0.01), respectively. In CaLu-3 cells, all XBB variants, including EG.5.1 and XBB.2.3, remained significantly lower in infectivity than D614G (p < 0.001) (**Fig. 1C**), as seen previously for Omicron-lineage variants^5-7,26,27^. EG.5.1 and its single mutations, XBB.1.5-G52H and XBB.1.5-F456L, exhibited comparable infectivity relative to parental XBB.1.5 (p > 0.05) (**Fig 1C**), with 1.4-fold (p > 0.05) and 1.3-fold increases (p < 0.05), respectively. XBB.2.3 also exhibited comparably infectivity relative to its parental XBB, with a 1.2-fold increase (p > 0.05) (**Fig 1C**). Overall, EG.5.1 and XBB.2.3 possess comparable infectivity to their parental XBB variants in ACE2 (HEK293T-ACE2) and CaLu-3 cells.

### EG.5.1 and XBB.2.3 exhibit comparable escape of neutralizing antibodies in bivalent vaccinated sera to other XBB-lineage subvariants

We next investigated escape of EG.5.1 and XBB.2.3 from neutralizing antibodies in serum samples collected from individuals that received at least 2 doses of monovalent mRNA vaccine and 1 dose of bivalent (wildtype + BA.4/5 spike) mRNA vaccine. These sera were collected from The Ohio State University Wexner Medical Center Health Care Workers (HCWs) at least three weeks post-booster administration. The neutralization assays were conducted with pseudotyped lentivirus as described previously^28^, and the cohort totaled 14 individuals (n = 14). Among these, 7 became positive during the Omicron wave, 3 tested positive prior to Omicron, and 4 were negative throughout. Sera were collected between 23 and 108 days after receiving a bivalent vaccination (median 66 days, **Table S1**). Consistent with previous results^5,7^, all XBB-lineage subvariants, including EG.5.1 and XBB.2.3, demonstrated marked reductions in antibody neutralization relative to D614G and BA.4/5^5,7^ (**Fig 2A-B**). EG.5.1 exhibited modestly decreased neutralization relative to XBB.1.5 (p > 0.05), which appeared to be driven by XBB.1.5-F456L mutation (**Fig 2A-B**). Notably, neutralizing antibody titers against EG.5.1 were markedly less than those against BA.4/5, with a 10-fold reduction (p < 0.01). Again, this phenotype was largely driven by the XBB.1.5-F456L mutation, which exhibited a 11.3-fold reduction in titer (p < 0.001) relative to BA.4/5 (**Fig 2A-B**). Furthermore, nAb titers of the 10 HCWs with breakthrough infection were much higher than those of the 4 HCWs without breakthrough infection (**Fig S1A**), indicating that breakthrough infection augments both the magnitude and breadth of nAbs. In contrast to EG.5.1, XBB.2.3 exhibited slightly increased neutralizing antibody titers relative to its parental XBB, with a 1.5-fold difference (p > 0.05). These titers were still lower than those against BA.4/5, with a 5.6-fold reduction (p < 0.001) (**Fig 2A-B**). Neither of the single mutations, XBB-D253G and XBB-P521S, exhibited distinct phenotypes in neutralization resistance from XBB.2.3 (**Fig 2A-B**). Overall, EG.5.1 and XBB.2.3 exhibit comparable escape of neutralizing antibodies in bivalent vaccinated sera to other XBB-lineage subvariants.

**Figure 2:**
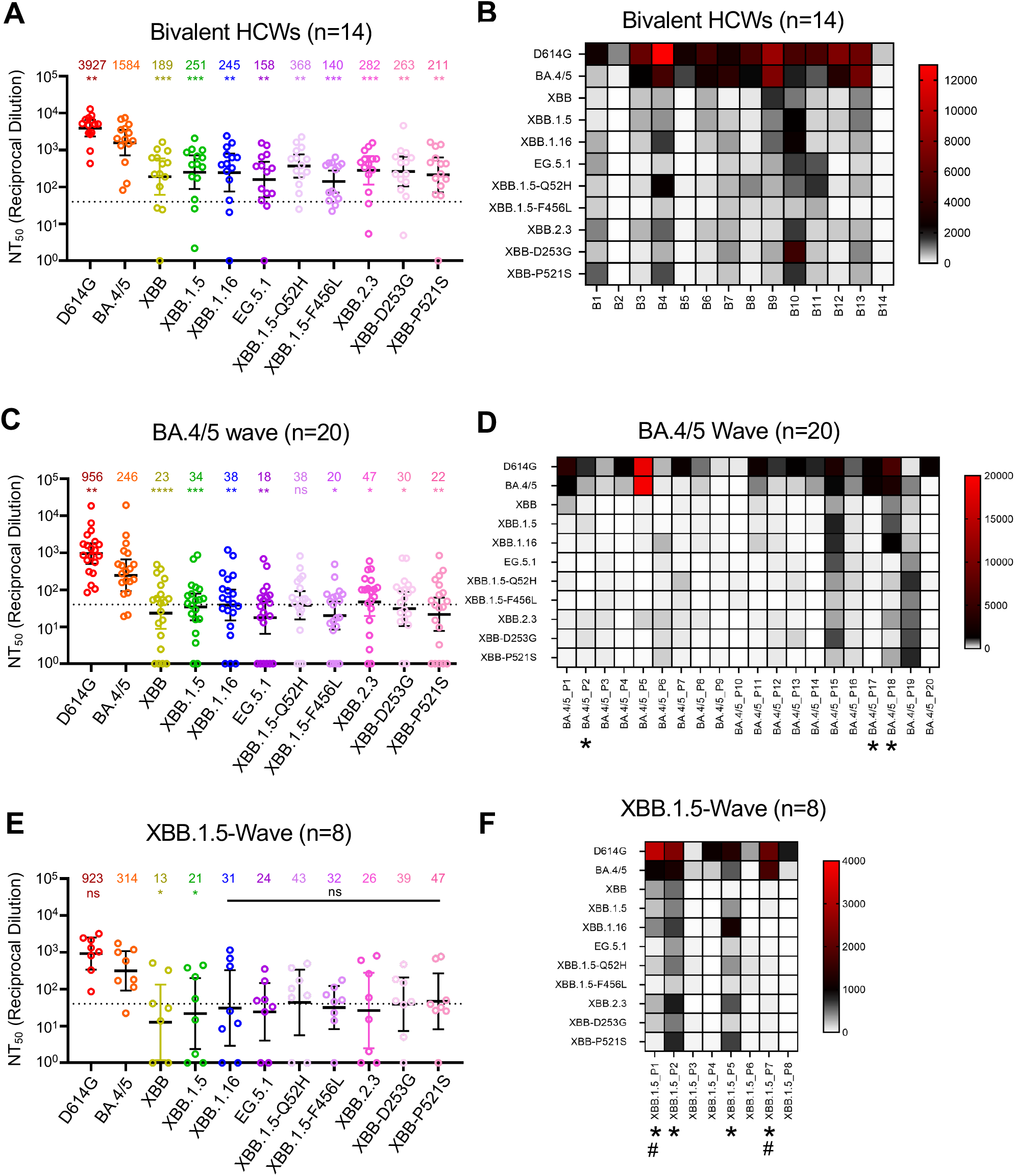
Neutralizing antibody titers against XBB.2.3 and EG.5.1 for bivalent vaccinees, BA.4/5-convalsecent cohort, and XBB.1.5-convalsecent cohort. Pseudotyped lentiviruses bearing each of the spikes of interest were used to perform virus neutralization assays with three cohorts of sera; **(A-B)** individuals that received at least two doses of monovalent mRNA vaccine and 1 dose of bivalent mRNA vaccine (n=14), **(C-D)** individuals that were infected during the BA.4/5-wave of COVID-19 in Columbus, OH (n=20); **(E-F)** individuals that were infected during the XBB.1.5-wave of COVID-19 in Columbus, OH (n=8). **(A, C, E)** Plots depict individual neutralizing antibody titers displayed as neutralization titers at 50% (NT_50_). Bars represent geometric means with 95% confidence intervals. Numbers on top of the plots represent the geometric means for each variant. Significance values are determined relative to BA.4/5, ancestor of these XBBs, using log10 transformed NT_50_ values to better approximate normality. **(B, D, F)** Heatmaps that depict the NT_50_ values for **(B)** the bivalent vaccinated cohort, **(D)** the BA.4/5-convalescent cohort, and **(F)** the XBB.1.5-convalscent cohort. Asterisks in **(D and F)** indicate the individuals who had received at least three doses of monovalent mRNA vaccine before infection. Hashtags in **(F)** indicate individuals that received at least 3 doses of monovalent mRNA vaccine and 1 dose of bivalent booster. p values are displayed as *p < 0.05, **p < 0.01, ***p < 0.001, ****p < 0.0001 and ns p > 0.05.

### EG.5.1 and XBB.2.3 markedly escape of neutralizing antibodies in BA.4/5-wave convalescent sera

The next cohort we tested were first responders and their household contacts who were infected during the BA.4/5-wave of COVID-19 in Columbus, OH (**Table S1**). Nasal swabs from these individuals confirmed COVID-19 positivity of 20 individuals (n = 20). Samples were sent for sequencing to determine the infecting variant; 4 individuals were infected with BA.4, 7 with BA.5, and 9 were undetermined but assumed to be infected with BA.4/5 based on the timing of collection when this variant was dominant in Columbus (July 2022 to late September 2022). In this cohort, 3 individuals had received 3 doses of either the Pfizer BioNTech BNT162b2 (n = 1) or Moderna mRNA-1272 (n = 2) vaccine, and 17 individuals were unvaccinated (**Table S1**). Similar to previous results^5,7^, all XBB-lineage subvariants exhibited marked escape of BA.4/5-wave convalescent sera, with all values under or around the limit of detection for the assay, i.e., 1:40^5,7^ (**Fig 2C-D, Fig S1B**). Both EG.5.1 and XBB.2.3 exhibited escape comparable to their parental variants (p > 0.05 for both) and had significant decreases in neutralizing antibody titer relative to BA.4/5, with reductions of 13.8-fold (p < 0.01) and 5.3-fold (p < 0.05), respectively (**Fig 2C-D, Fig S1B**).

### XBB.1.5-wave convalescent sera do not efficiently neutralize EG.5.1 and XBB.2.3

The third cohort we tested were 8 individuals from Columbus, OH who were infected during the XBB.1.5-wave (**Table S1**). Nasal swabs were all confirmed to be COVID-19 positive, with XBB.1.5 variant confirmed in 7, the remaining presumptive XBB based on collection date. Escape of neutralizing antibodies by XBB-lineage subvariants was comparable to the BA.4/5-convalsecent cohort, with all titers again near or below the limit of detection (**Fig 2E-F**). EG.5.1 had comparable titers relative to its parental XBB.1.5 (p > 0.05), exhibiting a 13.4-fold decrease relative to BA.4/5 (p < 0.05) (**Fig 2C-D, 2E-F**). XBB.2.3 exhibited comparable neutralizing antibody titers with its ancestor XBB (p > 0.05), but lower titers than BA.4/5 with an 8.8-fold decrease (p > 0.05) (**Fig 2C-D, 2E-F**). Notably, 3 patients, especially P2 and P5, and to a lesser extent P1, exhibited higher titers against XBB variants including EG.5.1 and XBB.2.3 (**Fig 2F, Fig S1C**). Not surprisingly, P2 and P5 had received 3 doses of monovalent mRNA vaccine (one with Moderna and another with Pfizer), and P1 was vaccinated with 3 doses of monovalent plus one dose of Moderna bivalent mRNA shots (**Table S1, Fig 2F, Fig S1C**). Interestingly, P7, who was a 64-year-old woman and had received 4 doses of monovalent and one dose of Moderna bivalent vaccines showed very high titers against D614G and BA.4/5, but barely detectable titers against all the XBB variants, including EG.5.1 and XBB.2.3 (**Table S1, Fig 2F)**. As would be expected, P3 and P6, who received 2 doses of monovalent of mRNA vaccine, as well as P4 and P8, whom were unvaccinated, showed low if any titers against XBB variants, although low titers against D614G/BA.4/5 were detected (**Table S1, Fig 2F, Fig S1C**).

### Monoclonal antibody S309 maintains neutralization efficacy against EG.5.1 and XBB.2

In addition to protection afforded through vaccination, monoclonal antibodies (mAb) represent a critical method to control COVID-19, especially in the early phase^29^. We thus tested S309, a class III monoclonal antibody, which has been shown previously to neutralize most Omicron-lineage subvariants, including XBB.1.5^7^. Here we found that S309 was still effective against both EG.5.1 and XBB.2.3, with inhibitory concentrations at 50% (IC_50_) of 2.7 μg/mL and 6.1 μg/mL, respectively (**Fig 3A-B**). EG.5.1 exhibited a comparable IC_50_ to other XBB variants, but a ∼3-fold increased IC_50_ compared to D614G (0.86 μg/mL); the IC_50_ values of XBB.1.5-Q52H and XBB.1.5-F456L were 2.1 and 2.2, respectively (**Fig 3A-B**). XBB.2.3 demonstrated a more marked increase in IC_50_ (6.1 μg/mL) compared to XBB (2.3 μg/mL), which appeared to be driven by the P521S mutation with an IC_50_ of 7.7 μg/mL (**Fig 3A-B**). Molecular modeling revealed that mutations in EG.5.1 and XBB.2.3 do not affect the ability of S309 to recognize the spikes. These mutations are located outside the epitope region of antibody S309. and are therefore less likely to influence the ability of S309 to recognize the spike protein (**Fig S2A**).

**Figure 3:**
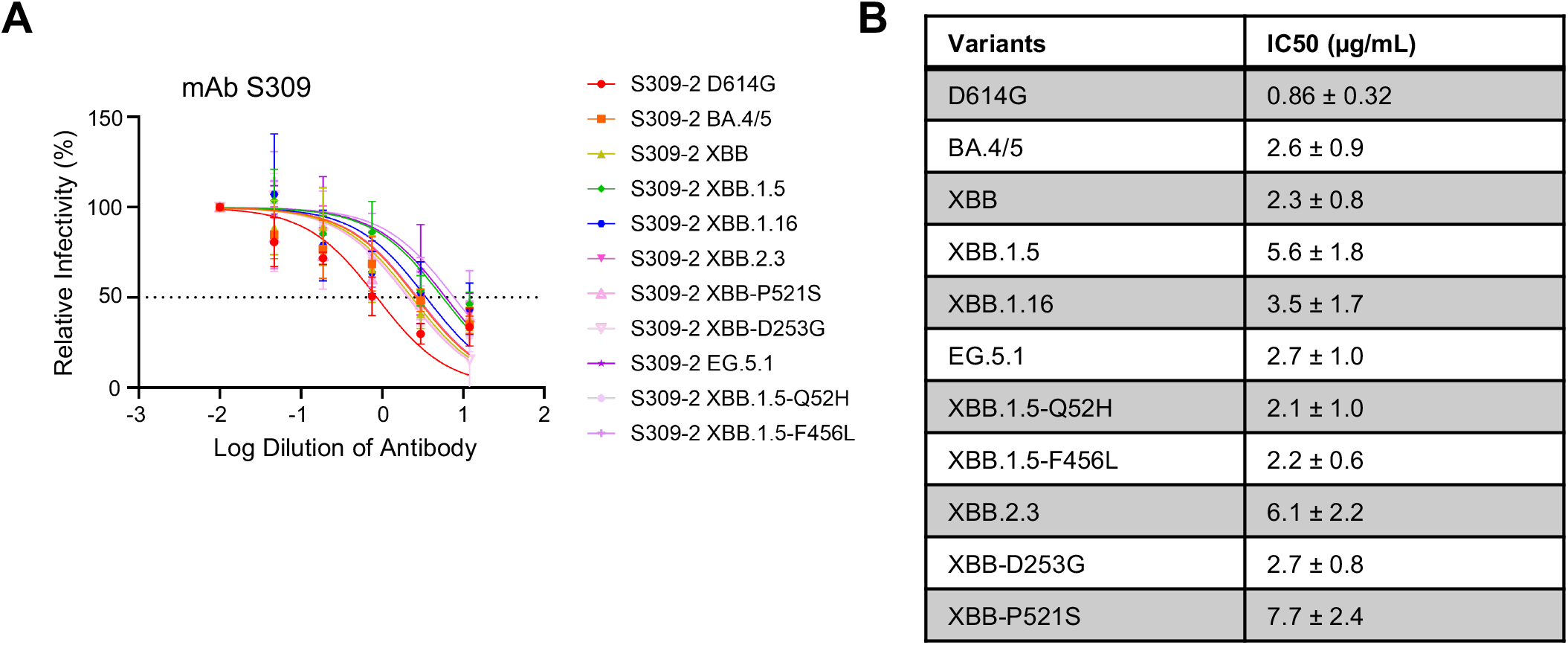
Neutralization of monoclonal antibody S309 against XBB.2.3 and EG.5.1. Pseudotyped lentiviruses bearing each of the spikes of interest were used in a virus neutralization assay with the class III monoclonal antibody S309. **(A)** Plot curve of S309 neutralization and **(B)** a table showing the calculated IC_50_ values best fit to the curve with the 95% confidence interval are depicted. The dashed line in **(A)** marks 50% relative infectivity.

### The fusion activities of XBB.2.3 and EG.5.1 spike are comparable to other XBB variants but lower than D614G

To determine the fusion activity of SARS-CoV-2 XBB spikes, we co-transfected HEK293T-ACE2 cells with GFP and the spike of interest and incubated the cells for 18 hours before imaging syncytia formation using fluorescence microscopy. We quantified the total area of fused cells using Leica X Applications Suite software implemented in Leica DMi8 microscope. Overall, EG.5.1 and XBB.2.3 showed a reduced fusogenicity relative to D614G, which is consistent with our previous results^5,7,27,30,31^. The fusion efficiency was comparable to other variants (**Fig 4A-B**), except XBB.1.16 (**Fig 4A-B**), which showed lower fusogenicity (*Faraone et al. Cell Reports* (in revision)). Surface expression levels of EG.5.1 and XBB.2.3 spikes on HEK293T producing pseudotyped lentiviruses were largely comparable, as shown by flow cytometry using an anti-S1 antibody (**Fig 4C-D**).

**Figure 4:**
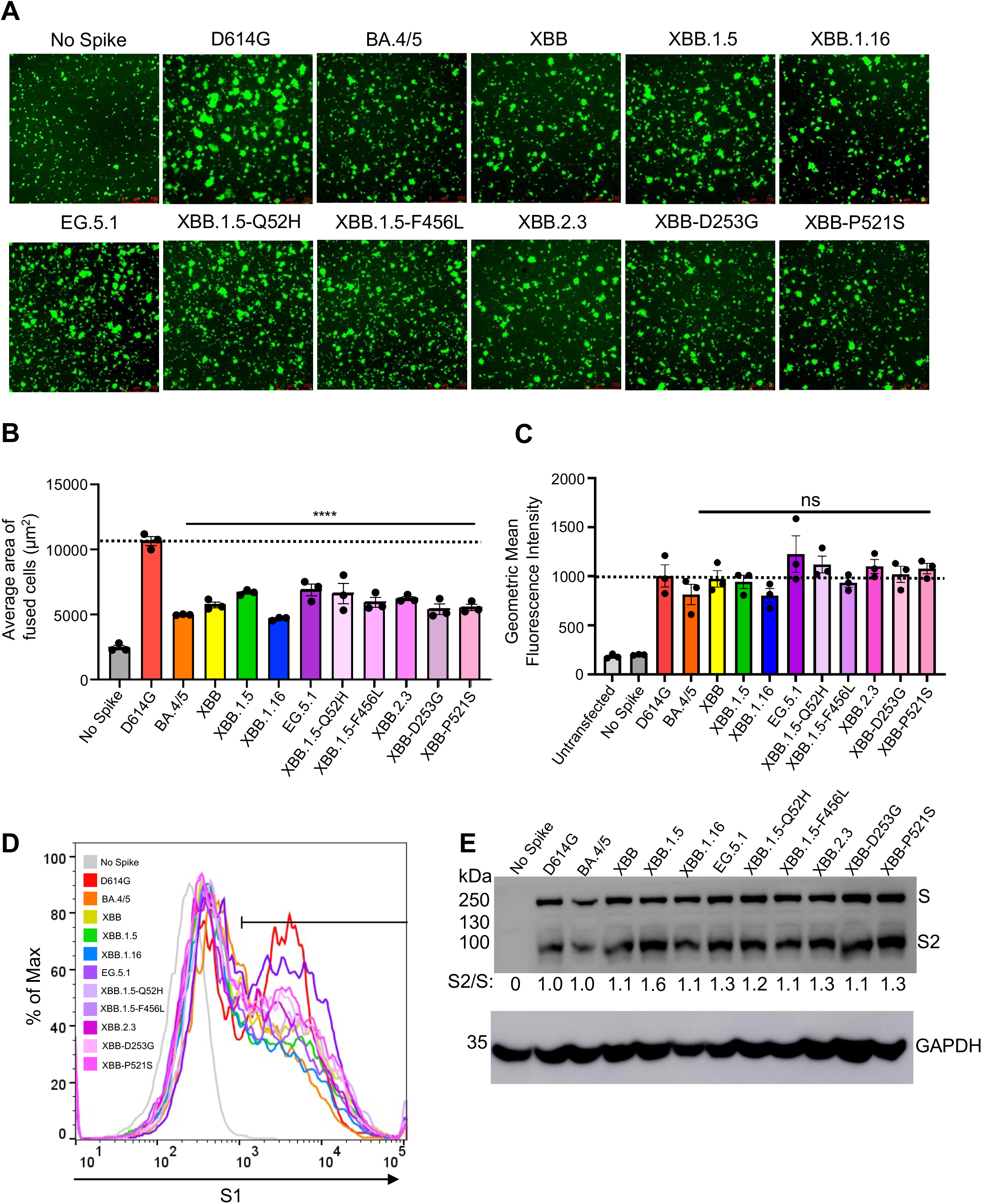
Fusogenicity, expression, and processing of XBB.2.3 and EG.5.1 spikes. **(A and B)** Fusogenicity of spikes were determined by co-transfecting HEK293T-ACE2 cells with GFP and the spike of interest and imaging the extent of fusion 18 hours post-transfection using fluorescence microscopy. **(A)** Representative images were selected and **(B)** average areas of fusion quantified for each spike. Bars represent means ± standard error, and dots represent three random areas for each replicate. Significance relative to D614G was determined using a one-way repeated measures ANOVA with Bonferroni’s multiple testing correction (n = 3). “No Spike” refers to the negative control which was transfected with GFP and empty pcDNA3.1 plasmid. **** p < 0.0001. **(C)** Expression of spike was determined by performing surface staining (anti-S1 polyclonal antibody) and flow cytometry on HEK293T cells used to produce pseudotyped lentiviruses. A triplicate was performed, and a representative overlaid histogram was selected and depicted in **(C). (D)** The processing of each spike was determined by lysing HEK293T cells transfected with spike of interest and performing western blotting. Blots were probed with anti-S2 and anti-GAPDH (loading control), respectively. Processing of spike was quantified using Image J to determine relative band intensities for full length spike versus S2 and a resulting S2/S ratio was calculated. Ratios are listed below each corresponding set of bands. Ratios were normalized to D614G (D614G=1.0).

We also investigated the processing of each spike into its S2 subunits in lysates of transfected HEK293T cells. We performed western blotting and probed with an anti-S2 polyclonal antibody to compare the ratios between S2 and full-length spike among the variant spikes tested. As shown **Fig 4E**, EG.5.1 and XBB.2.3 exhibited efficiencies of spike processing comparable to other XBB variants, the levels of which were generally higher than that of D614G.

### Decreased antigenic distance in bivalent vaccinated relative to convalescent cohorts

To better understand how antigenicity varies between variants, we conducted antigenic mapping analysis on the three sets of neutralization titers presented above^32^. The method uses multidimensional scaling on log2 transformed binding assay results to plot individual points for antigens and antibodies in Euclidean space^32^. The spaces between the different points directly translate from fold changes in neutralization titers, allowing for visualization of the antigenic differences between the variant spikes. The points are plotted using “antigenic distance units” (AU), with one AU being equivalent to a 2-fold change in neutralizing antibody titer^13,32^. In all cohorts, D614G and BA.4/5 clustered together while XBB variants were more antigenically distinct, sitting around 4.0-5.5 AU away from D614G, translating to a 16∼45-fold drop in overall neutralizing antibody titer (**Fig 5A-C, Fig 2**). Antigenic distance between all variants was overall slightly smaller for the bivalent relative to the convalescent cohorts (**Fig 5A-C**), suggesting a broader neutralization induced by the bivalent vaccine. XBB.2.3 consistently clustered with XBB.1.16, whereas EG.5.1 appeared more antigenically distant from the other XBB-lineage variants (**Fig 5A-C**). This phenotype was more pronounced in the XBB.1.5-wave cohort (**Fig 5C**). Overall, XBB-lineage variants are notably distinct antigenically from earlier variants D614G and BA.4/5, but this is somewhat minimized upon bivalent vaccination.

**Figure 5:**
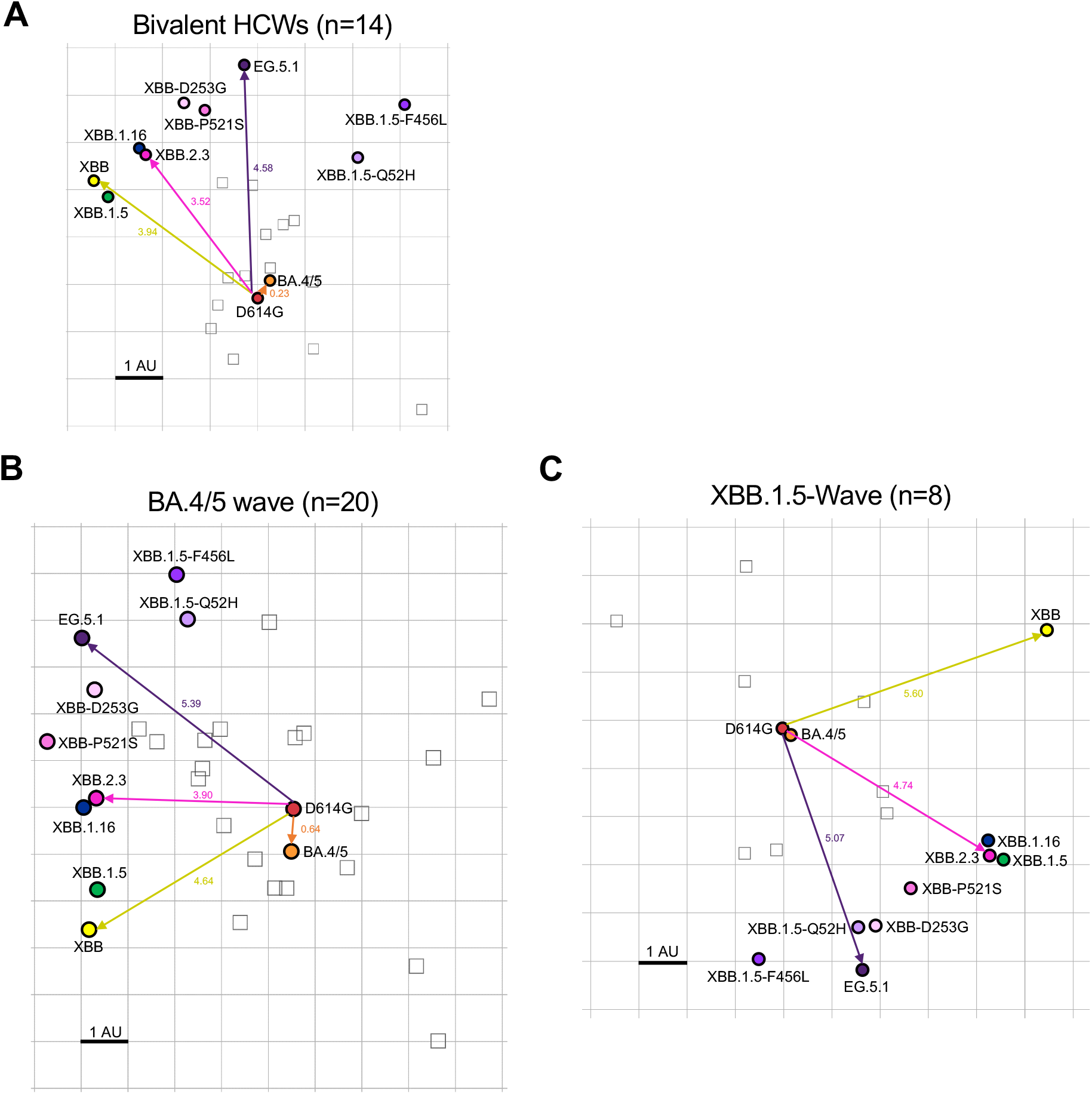
Antigenic mapping of neutralization titers for bivalent vaccinated, BA.4/5-wave infected, and XBB.1.5-wave infected cohorts (associated with Fig 2). The Racmacs program (1.1.35) was used to generate antigenic maps for neutralization titers from **(A)** the bivalent vaccinated, **(B)** the BA.4/5 wave infected, and **(C)** the XBB.1.5-wave infected cohorts. Circles represent the variants and squares represent the individual sera samples. Arrows between D614G and selected variants are labeled with the distance between those variants in antigenic units (AU). One square on the grid represents one antigenic unit squared.

## Discussion

SARS-CoV-2 continues to evolve rapidly, presenting an ever-increasing challenge to vaccination efforts. As expected, new XBB-lineage subvariants EG.5.1 and XBB.2.3, especially EG.5.1, remain highly immune evasive, which likely contributes to the recent increase of COVID cases and hospitalization^23,24^. Though bivalent vaccination continues to protect better than the monovalent vaccine and natural infection, neutralization titers are markedly low against all XBB variants, particularly the newly emerged EG.5.1, in comparison to D614G and BA.4/5, as seen previously for XBB variants^3,5,7,9,12,14,16^. Neutralizing antibody titers stimulated by infection with either BA.4/5 or XBB.1.5 are minimal, with average neutralization titers against XBB variants clustering around the limit of detection for the assay, which is consistent with another study^33^. We found that the XBB.1.5-F456L mutation, rather than the XBB.1.5-Q52H mutation, drives the enhanced neutralization escape compared to XBB.1.5. Molecular modeling indicates that XBB.1.5-F456L likely decreases spike binding to class 1 SARS-CoV-2 monoclonal antibodies, such as antibody S2E12, but does not appear to impact spike binding to S309, a class 3 monoclonal antibody (**Fig S2A-B**), a finding that is consistent the several recent publications^33-35^. Together, these studies underscore the need for close surveillance of variants and newly formulated vaccines against SARS-CoV-2.

Notably, in our study, bivalent-vaccinated neutralizing antibody titers against BA.4/5 were distinguishably lower than D614G despite BA.4/5 spike being included in the vaccine formulation (**Fig 2A-B**). This suggests that the antibody response is still largely targeting D614G, hence providing evidence for immune imprinting induced by the monovalent doses of mRNA vaccines^20-22,34,36^. Many mutations have been acquired by the virus during its evolution from BA.4/5 through the various XBB variants^37^. Notably, neutralizing antibody titers for the bivalent cohort against XBB variants remain significantly lower than D614G and BA.4/5 (**Fig 2A-B)**. Consistently, antigenic mapping demonstrates that XBB variants are quite antigenically distinct from D614G and BA.4/5 for all cohorts tested, especially EG.5.1 (**Fig 2, Fig 5**). Importantly, the distinct antigenic phenotype of XBB and other Omicron subvariants has been corroborated by other studies using antigenic cartography analysis^21,22,38^.

We observed that the antigenic distance between all variants was smaller overall for the bivalent vaccination cohort, the majority of which had breakthrough infection, relative to the convalescent cohorts (**Fig 5, Table S1**). Two of 4 vaccinated individuals infected with XBB.1.5, i. e., P2 and P5, did exhibit the broadest neutralizing antibody titers among the cohort (**Fig 2E-F**), suggesting that vaccines containing XBB.1.5 and related spikes, such as XBB.1.16, EG.5.1, will likely overcome immune imprinting and offer broader protection against XBB-lineage subvariants. This finding suggests that the bivalent vaccine/breakthrough combination increases coverage of immune responses against newer SARS-CoV-2 variants, as has been suggested previously by another group^20^ (**Fig S2A, Fig 5**). Fortunately, newly formulated mRNA vaccines containing XBB.1.5 spike have been submitted by Pfizer and Moderna to the FDA and are expected to rollout in September^39-41^.

We did not find dramatic changes in infectivity of EG.5.1 and XBB.2.3 compared to previous XBB variants in either 293T-ACE2 or CaLu-3 cells (**Fig 1**). EG.5.1 had a modest increase in 293T-ACE2 but this was not observed in the more biologically relevant lung airway epithelial cell line CaLu-3. Importantly, similar to Omicron variants BA.4/5 and XBB variants, EG.5.1 and XBB.2.3 retain lower infectivity in CaLu-3 cells relative to D614G, easing concerns of potentially increased pathogenesis in the lung. Furthermore, the fusogenicity of EG.5.1 and XBB.2.3 are similar to other XBB variants, which is much lower than D614G (**Fig 4**). In this regard, molecular modeling reveals that the XBB.1.5-F456L mutation may reduce spike binding to ACE2 (**Fig S2C**). Specifically, the change from phenylalanine to leucine decreases the side chain size and increases the distance between the receptor-binding domain (RBD) and ACE2 residues, resulting in a reduction of hydrophobic interactions at this specific position. Hence, the affinity between viral RBD and the ACE2 receptor is likely diminished. Overall, while we did not find in vitro evidence to support that the newly emerged XBB variants, including EG.5.1, have enhanced pathogenic potential that could explain a possible growth advantage in circulation around the globe^24,33^, in vivo assays and clinical studies are needed to address this important issue.

Overall, our study provides important support for new vaccine formulations in our quest for continued control of the COVID-19 pandemic, underscored by the marked immune evasion of XBB variants^5,7,9,12,14,16,17,33,42^ and the role of immune imprinting in these phenotypes^20-22,38^. Specifically, removal of the wildtype spike from mRNA vaccines and inclusion of XBB-lineage variant spikes must be considered. The continued surveillance of new variants is essential to inform decisions around vaccination against SARS-CoV-2 and treatment of COVID-19.

### Limitations of the Study

Pseudotyped virus was used throughout the study in place of live authentic viruses. We have previously validated our neutralization assay alongside live virus^28^, and we believe the timeliness of the work justifies the use of pseudotyped virus over live virus. Pseudotyped virus also provides critical advantage for investigating the role of specific spike variants in neutralization, membrane fusion and infectivity in a more controlled manner. Our cohort sizes for the neutralization assays were small, particularly the XBB.1.5-wave cohort, because of the difficulty in recruiting as result of the decreased COVID-19 testing. However, we believe our findings are still valid and significant given that other groups have published such work with comparable cohorts and similar methods^16,34^, and that our findings for XBB.1.5-wave individuals corroborate results from another group^33^. The sample collection time after vaccination or infection also varies widely in our cohorts due to the clinical arrangements, which could have impacted the nAb titers. Overall, we feel as though these limitations are outweighed by the timeliness of the work and the importance of the continued characterization of variants to maintain control of the COVID-19 pandemic.

## STAR Methods

### Resource availability

#### Lead contact

The lead contact Dr. Shan-Lu Liu can be reached at with any questions or requests for materials.

#### Materials availability

Materials can be requested by emailing the lead contact.

### Experimental model and subject details

#### Vaccinated and convalescent cohorts

Three cohorts of serum were collected and used to determine neutralizing antibody titers against selected SARS-CoV-2 variants. The first were health care workers (HCWs) working at the Ohio State Wexner Medical Center that received at least 2 doses of monovalent mRNA vaccine and 1 dose of bivalent mRNA vaccine. Samples were collected under the approved IRB protocols 2020H0228, 2020H0527, and 2017H0292. This cohort totaled 14 individuals, i.e., 8 males and 6 females. Among these, 12 individuals received 3 doses of monovalent vaccine (Pfizer BioNTech BNT162b2 or Moderna mRNA-1273) and 1 dose of bivalent vaccine (Pfizer); 1 individual had 2 doses of monovalent vaccine (Pfizer) and 1 dose bivalent (Pfizer), and the final individual had 4 doses of monovalent vaccine (Pfizer) and 1 dose of bivalent vaccine (Pfizer). Sample collections ranged from 23-108 days post-administration of booster dose and the range of ages was 25-48 (median 36).

The second cohort were first responders and household contacts based in Columbus, OH that were infected with SARS-CoV-2 during the BA.4/5-wave of infection in Columbus. Samples were collected under approved IRB protocols 2020H0527, 2020H0531, and 2020H0240. This cohort totaled 20 individuals. For each, nasal swabs were used to confirm positive infection with the virus, and were sequenced. 4 individuals were confirmed to be infected with BA.4, 7 with BA.5, and the remaining 9 were undetermined but assumed to be infected with BA.4/5 due to timing of collection during when the variant was dominant in Columbus (July 2022 to late September 2022). 3 individuals in this cohort were vaccinated with 3 doses of monovalent vaccine (1 Pfizer and 2 Moderna). The age range of this cohort was 27-58 (median 44), and it included 4 male, 15 female, and 1 unknown individuals.

The final cohort were first responders that were infected during the XBB.1.5 wave in Columbus, Ohio (Early February 2023 through July 2023). Samples were collected under IRB protocols 2020H0527, 2020H0531, and 2020H0240. The cohort totaled 8 individuals (n=8). Like the BA.4/5-wave samples, nasal swabs were performed on each member of the cohort and the samples were sent for sequencing. Seven samples were confirmed to be XBB.1.5 by COVID-Seq Artic v4 sequencing and typing with Dragen COVID Lineage, with Pangolin plug-in (Illumina), with one presumptive XBB.1 based on date of collection. Several showed private or regional variations in spike (e.g. T284I and L513F). 5 individuals were vaccinated with at least two doses of monovalent mRNA vaccine while 4 were not vaccinated. Of the vaccinated members of the cohort, 1 received two doses of monovalent Moderna mRNA vaccine, 2 individuals received 3 doses of monovalent vaccine (1 Pfizer, 1 Moderna), 1 individual received 3 doses of monovalent vaccine and 1 dose of bivalent (all Moderna), and the last vaccinated person received 4 doses of monovalent vaccine with 1 dose of bivalent (Moderna monovalent, Pfizer bivalent). The range of ages was 38-64 (median 53), and the cohort had 5 male and 3 female individuals. Full details of each cohort can be found in **Table S1**.

#### Cell lines

Cell lines used for this study included human embryonic kidney line HEK293T (ATCC CRL-11268, RRID: CVCL_1926), HEK293T expressing human ACE2 (HEK293T-ACE2) (BEI NR-52511, RRID: CVCL_A7UK), and human adenocarcinoma lung epithelial line CaLu-3 (RRID: CVCL_0609). HEK293T and HEK293T-ACE2 cells were maintained DMEM (Gibco, 11965-092) supplemented with 10% fetal bovine serum (Sigma, F1051) and 0.5% penicillin-streptomycin (HyClone, SV30010). CaLu-3 cells were maintained in EMEM supplemented the same way. To split, cells were initially washed with phosphate-buffered saline (Sigma, D5652-10X1L) then incubated in 0.05% trypsin + 0.53mM EDTA (Corning, 25-052-CI) until complete detachment. Cells were kept at 37C and 5.0% CO2.

### Method Details

#### Plasmids

All spike plasmids are in the backbone of pcDNA3.1 with restriction sites BamHI and KpnI and FLAG tags at the N- and C-termini of spike. D614G, BA.4/5, and XBB plasmids were cloned by GenScript using restriction enzyme cloning (Piscataway, NJ). XBB.1.5, XBB.1.16, XBB-D253G, XBB-P521S, XBB.2.3, XBB.1.5-Q52H, XBB.1.5-F456L, and EG.5.1 plasmids were generated in house through site-directed mutagenesis. Throughout, the “No Spike” control refers to empty pcDNA3.1 plasmid backbone used in place of spike plasmid. The lentiviral vector used is a HIV-1, pNL4-3 vector with an Env deletion and intronic secreted *Gaussia* luciferase reporter (inGluc).

#### Pseudotyped lentivirus production and infectivity

Pseudotyped lentiviral vectors were produced by co-transfecting HEK293T cells in a 2:1 ratio with pNL4-3 inGluc and the spike plasmid of interest. Transfections throughout are polyethyleneimine transfections using the Transporter 5 transfection reagent (Polysciences). Pseudotyped virus was collected 48 and 72 hours post-transfection and stored at -80C. To measure infectivity, 100uL of virus was used to infect HEK293T-ACE2 cells. 300uL was used to infect CaLu-3 cells and cells were spun at 1,650 x g for 45 minutes to mediate attachment. Luciferase measurements were taken as a readout of infectivity at 48, 72, and 96 hours. Measurements were collected by taking 20uL of infected cell media and combining it with 20uL luciferase substrate (0.1 M Tris pH 7.4, 0.3 M sodium ascorbate, 10 μM coelenterazine) and immediately reading on a BioTek Cytation plate reader. Plots for 48 hours and 120 hours are displayed in Fig 1 for HEK293T-ACE2 and CaLu-3, respectively.

#### Virus neutralization assay

Sera from the cohorts of interest was first serially diluted four-fold with a starting dilution of 1:40 (final dilutions 1:40, 1:160, 1:640, 1:2560, 1:10240, and no serum as a control). mAb S309 was diluted 4-fold from 12 μg/ml (12, 3, 0.75, 0.1875, 0.046875μg/ml, no antibody control). Pseudotyped virus was thawed and diluted based on infectivity results to normalize readouts. 100uL of each diluted virus was then added onto serum samples. The virus and sera mixture were incubated for 1 hour at 37C. This mixture was then used to infect HEK293T-ACE2 cells. Luciferase readout was collected as described above at 48 and 72 hours post-infection. NT_50_ values were determined through least-squares fit non-linear regression with a normalized response (no serum control) in GraphPad Prism 9 (San Diego, CA).

#### Syncytia formation

HEK293T-ACE2 cells were co-transfected with GFP and the spike of interest. Cells were imaged 18 hours post-transfection using a Leica DMi8 fluorescence microscope. Average area of fused cells was determined using the Leica X Applications Suite software that outlines edges of syncytia and calculates the area within. Three images were randomly taken for each variant. Scale bars represent 150 μM and one representative image was selected for presentation.

#### S protein surface expression

Seventy-two hours post transfection, HEK293T cells used to produce lentivirus were washed in PBS and incubated in PBS+5mM EDTA for 10 minutes to detach. Approximately 1x10^6 cells were taken for analysis of spike surface expression via flow cytometry. These cells were fixed in 3.7% formaldehyde for 10 minutes and room temperature. Cells were stained with 1:200 anti-S1 polyclonal antibody (Sino Biological, 40591-T62; RRID: AB_2893171) for 1.5 hours and washed three times in PBS+2% FBS. Cells were then stained with secondary antibody 1:200 anti-Rabbit-IgG-FITC (Sigma, F9887, RRID: AB_259816) and washed three times more. Flow cytometry was performed on a LifeTechnologies Attune NxT flow cytometer. Data processing was performed using FlowJo v10.9.1 (Ashland, OR).

#### S protein processing

HEK293T cells transfected with spike of interest were lysed in 300uL RIPA+PI+PMSF (RIPA: 50mM Tris pH 7.5, 150 mM NaCl, 1 mM EDTA, Nonidet P-40, 0.1% SDS, PI+PMSF: Sigma, P8340) for 40 minutes on ice. Lysate was harvested and used for western blotting. Samples were run on a 10% acrylamide SDS-PAGE gel and transferred to a PVDF membrane. Blots were probed with anti-S2 (Sino Biological, 40590; RRID:AB_2857932) and anti-GAPDH as a loading control (Santa Cruz, Cat# sc-47724, RRID: AB_627678). Secondary antibodies included anti-Rabbit-IgG-FITC (Sigma, A9169; RRID:AB_258434) and anti-Mouse-IgG-FITC (Sigma, Cat# A5278, RRID: AB_258232). Blots were imaged using Immobolin Crescendo Western HRP substrate (Millipore, WBLUR0500) and exposed on a GE Amersham Imager 600. Quantification of band intensity was determined using ImageJ (NIH, Bethesda, MD).

#### Antigenic mapping

Antigenic maps were generated using Racmacs (v1.1.35) (https://github.com/acorg/Racmacs/tree/master). This method is based on a study conducted by Smith and colleagues to determine the antigenic distances between different influenza strains based on agglutination neutralization assays^32^. Briefly, raw neutralization titers were converted into a table with sera samples as the columns and viruses as the rows. This table was then imported into the *Racmacs* program using R (Vienna, Austria) and instructions in the documentation section for the program were followed. *Racmacs* takes the titer table and converts it to a distance table by performing a log2 conversion and then calculating the distance between each antigen for each serum sample. Multidimensional scaling is then performed on the distance table to generate the map. Optimization settings were kept on default (2 dimensions, 500 optimizations, minimum column basis “none”). Maps were saved from the “view” panel and labeled using Microsoft Office PowerPoint. Arrows drawn in PowerPoint were used to calculate the distance between two points with the scale bar for “1 AU” being used to normalize this value. 1 AU is equivalent to a 2-fold change in neutralizing antibody titer^13,32^.

#### Structural Modeling and Analysis

We conducted structural modeling of EG.5.1 spike proteins bound to either the ACE2 receptor or neutralizing antibodies. This modeling was carried out using the SWISS-MODEL server, employing existing X-ray crystallography or cryo-EM structures from published sources as templates (PDB: 7K8Z, 8DT3, 7L7D, 7XB0, 7XCK, 7YAD, 7R6X). Molecular interactions involving EG.5.1 mutants were carefully examined, and these interactions were visually presented using PyMOL.

#### Quantification and statistical analysis

Statistical analyses were performed using GraphPad Prism 9. Error bars in **(Fig 1B-C)** and **(Fig 4C)** represent means with standard error. Comparisons between the viruses in **(Fig 1B-C)** and **(Fig 4C)** were made using a one-way ANOVA with Bonferroni post-test. Both experiments (infectivity and surface expression) were done in triplicate. Neutralization titers were determined using least-squares non-linear regression. Error bars in **(Fig 2A, C, and E)** represent geometric means with 95% confidence intervals. Comparisons between the viruses in **(Fig 2A, C, and E**) were made using a repeated measures one-way ANOVA with Bonferroni post-test. These comparisons were conducted using log10 transformed NT_50_ values to better approximate normality. Bars in **(Fig 3)** represent best fit values for IC_50_ ± 95% confidence interval (n=1). Significance analysis in (**Fig 4**) was performed using a one-way repeated measures ANOVA with Bonferroni’s multiple testing correction.

## Data and code availability

Data can be requested from the lead contact. This paper does not report original code.

## Acknowledgements

We thank the Clinical Research Center/Center for Clinical Research Management of The Ohio State University Wexner Medical Center and The Ohio State University College of Medicine in Columbus, Ohio, specifically J. Brandon Massengill, Francesca Madiai, Dina McGowan, Breona Edwards, Evan Long, and Trina Wemlinger, for logistics, collection, and processing of samples. We thank Tongqing Zhou at NIH for providing the S309 monoclonal antibody. In addition, we thank Sarah Karow, Madison So, Preston So, Daniela Farkas, and Finny Johns in the clinical trials team of The Ohio State University for sample collection and other supports. We thank Ashish R. Panchal, Soledad Fernandez, Mirela Anghelina, and Patrick Stevens for their assistance in providing the sample information of the first responders and their household contacts. We thank Peng Ru and Lauren Masters for sequencing and Xiaokang Pan for bioinformatic analysis. S.-L.L., D. J., R.J.G., L.J.S. and E.M.O. were supported by the National Cancer Institute of the NIH under award no. U54CA260582. The content is solely the responsibility of the authors and does not necessarily represent the official views of the National Institutes of Health. This work was also supported by a fund provided by an anonymous private donor to OSU. K.X. was supported by The Ohio State University Comprehensive Cancer Center, a Path to K grant through the Ohio State University Center for Clinical & Translational Science. R.J.G. was additionally supported by the Robert J. Anthony Fund for Cardiovascular Research and the JB Cardiovascular Research Fund, and L.J.S. was partially supported by NIH R01 HD095881.

## Author contributions

S.-L.L. conceived and directed the project. R.J.G led the clinical study/experimental design and implementation. J.N.F performed neutralization and infectivity assays, and P.Q. performed syncytia formation and spike processing, N.G. performed mutagenesis to generate new variants. P.Q and J.N.F. performed data processing and analyses. D.J. led SARS-CoV-2 variant genotyping and DNA sequencing analyses. C.C., and R.J.G. provided clinical samples and related information. K.X. performed molecular modeling and participated in discussion. J.N.F., P.Q., and S.-L.L. wrote the paper. Y.-M.Z, L.J.S., and E.M.O. provided insightful discussion and revision of the manuscript.

## References

1. Wang, Q., Guo, Y., Iketani, S., Nair, M.S., Li, Z., Mohri, H., Wang, M., Yu, J., Bowen, A.D., Chang, J.Y., et al. (2022). Antibody evasion by SARS-CoV-2 Omicron subvariants BA.2.12.1, BA.4 and BA.5. Nature 608, 603–608. 10.1038/s41586-022-05053-w.

2. Planas, D., Saunders, N., Maes, P., Guivel-Benhassine, F., Planchais, C., Buchrieser, J., Bolland, W.H., Porrot, F., Staropoli, I., Lemoine, F., et al. (2022). Considerable escape of SARS-CoV-2 Omicron to antibody neutralization. Nature 602, 671–675. 10.1038/s41586-021-04389-z.

3. Miller, J., Hachmann, N.P., Collier, A.Y., Lasrado, N., Mazurek, C.R., Patio, R.C., Powers, O., Surve, N., Theiler, J., Korber, B., and Barouch, D.H. (2023). Substantial Neutralization Escape by SARS-CoV-2 Omicron Variants BQ.1.1 and XBB.1. N Engl J Med 388, 662–664. 10.1056/NEJMc2214314.

4. Hachmann, N.P., Miller, J., Collier, A.Y., Ventura, J.D., Yu, J., Rowe, M., Bondzie, E.A., Powers, O., Surve, N., Hall, K., and Barouch, D.H. (2022). Neutralization Escape by SARS-CoV-2 Omicron Subvariants BA.2.12.1, BA.4, and BA.5. N Engl J Med 387, 86–88. 10.1056/NEJMc2206576.

5. Faraone, J.N., Qu, P., Evans, J.P., Zheng, Y.-M., Carlin, C., Anghelina, M., Stevens, P., Fernandez, S., Jones, D., Lozanski, G., et al. (2023). Neutralization escape of Omicron XBB, BR.2, and BA.2.3.20 subvariants. Cell Reports Medicine 4, 101049.

6. Evans, J.P., Zeng, C., Qu, P., Faraone, J., Zheng, Y.-M., Carlin, C., Bednash, J.S., Zhou, T., Lozanski, G., Mallampalli, R., et al. (2022). Neutralization of SARS-CoV-2 Omicron sub-lineages BA.1, BA.1.1, and BA.2. Cell host & microbe 30, 1093–1102.e1093. 10.1016/j.chom.2022.04.014.

7. Qu, P., Faraone, J.N., Evans, J.P., Zheng, Y.-M., Carlin, C., Anghelina, M., Stevens, P., Fernandez, S., Jones, D., Panchal, A.R., et al. (2023). Enhanced evasion of neutralizing antibody response by Omicron XBB.1.5, CH.1.1, and CA.3.1 variants. Cell Reports 42, 112443.

8. Cao, Y., Yisimayi, A., Jian, F., Song, W., Xiao, T., Wang, L., Du, S., Wang, J., Li, Q., Chen, X., et al. (2022). BA.2.12.1, BA.4 and BA.5 escape antibodies elicited by Omicron infection. Nature 608, 593–602. 10.1038/s41586-022-04980-y.

9. Zou, J., Kurhade, C., Patel, S., Kitchin, N., Tompkins, K., Cutler, M., Cooper, D., Yang, Q., Cai, H., Muik, A., et al. (2023). Neutralization of BA.4-BA.5, BA.4.6, BA.2.75.2, BQ.1.1, and XBB.1 with Bivalent Vaccine. N Engl J Med 388, 854–857. 10.1056/NEJMc2214916.

10. Zhang, X., Chen, L.L., Ip, J.D., Chan, W.M., Hung, I.F., Yuen, K.Y., Li, X., and To, K.K. (2022). Omicron sublineage recombinant XBB evades neutralising antibodies in recipients of BNT162b2 or CoronaVac vaccines. Lancet Microbe. 10.1016/s2666-5247(22)00335-4.

11. Yue, C., Song, W., Wang, L., Jian, F., Chen, X., Gao, F., Shen, Z., Wang, Y., Wang, X., and Cao, Y. (2023). Enhanced transmissibility of XBB.1.5 is contributed by both strong ACE2 binding and antibody evasion. bioRxiv, 2023.2001.2003.522427. 10.1101/2023.01.03.522427.

12. Yamasoba, D., Uriu, K., Plianchaisuk, A., Kosugi, Y., Pan, L., Zahradnik, J., Consortium, T.G.t.P.J., Ito, J., and Sato, K. (2023). Virological characteristics of the SARS-CoV-2 Omicron XBB.1.16 variant. bioRxiv, 2023.2004.2006.535883. 10.1101/2023.04.06.535883.

13. Wang, Q., Iketani, S., Li, Z., Liu, L., Guo, Y., Huang, Y., Bowen, A.D., Liu, M., Wang, M., Yu, J., et al. (2023). Alarming antibody evasion properties of rising SARS-CoV-2 BQ and XBB subvariants. Cell 186, 279–286.e278. 10.1016/j.cell.2022.12.018.

14. Uraki, R., Ito, M., Furusawa, Y., Yamayoshi, S., Iwatsuki-Horimoto, K., Adachi, E., Saito, M., Koga, M., Tsutsumi, T., Yamamoto, S., et al. (2023). Humoral immune evasion of the omicron subvariants BQ.1.1 and XBB. Lancet Infect Dis 23, 30–32. 10.1016/s1473-3099(22)00816-7.

15. Tamura, T., Ito, J., Uriu, K., Zahradnik, J., Kida, I., Anraku, Y., Nasser, H., Shofa, M., Oda, Y., Lytras, S., et al. (2023). Virological characteristics of the SARS-CoV-2 XBB variant derived from recombination of two Omicron subvariants. Nat Commun 14, 2800. 10.1038/s41467-023-38435-3.

16. Kurhade, C., Zou, J., Xia, H., Liu, M., Chang, H.C., Ren, P., Xie, X., and Shi, P.Y. (2022). Low neutralization of SARS-CoV-2 Omicron BA.2.75.2, BQ.1.1 and XBB.1 by parental mRNA vaccine or a BA.5 bivalent booster. Nat Med. 10.1038/s41591-022-02162-x.

17. He, Q., Wu, L., Xu, Z., Wang, X., Xie, Y., Chai, Y., Zheng, A., Zhou, J., Qiao, S., Huang, M., et al. (2023). An updated atlas of antibody evasion by SARS-CoV-2 Omicron sub-variants including BQ.1.1 and XBB. Cell Rep Med 4, 100991. 10.1016/j.xcrm.2023.100991.

18. Davis-Gardner, M.E., Lai, L., Wali, B., Samaha, H., Solis, D., Lee, M., Porter-Morrison, A., Hentenaar, I.T., Yamamoto, F., Godbole, S., et al. (2022). Neutralization against BA.2.75.2, BQ.1.1, and XBB from mRNA Bivalent Booster. New England Journal of Medicine 388, 183–185. 10.1056/NEJMc2214293.

19. Recommendation for the 2023-2024 Formula of COVID-19 vaccines in the U.S. (2023).

20. Yisimayi, A., Song, W., Wang, J., Jian, F., Yu, Y., Chen, X., Xu, Y., Yang, S., Niu, X., Xiao, T., et al. (2023). Repeated Omicron infection alleviates SARS-CoV-2 immune imprinting. bioRxiv, 2023.2005.2001.538516. 10.1101/2023.05.01.538516.

21. Wang, Q., Guo, Y., Tam, A.R., Valdez, R., Gordon, A., Liu, L., and Ho, D.D. (2023). Deep immunological imprinting due to the ancestral spike in the current bivalent COVID-19 vaccine. bioRxiv, 2023.2005.2003.539268. 10.1101/2023.05.03.539268.

22. Cao, Y., Jian, F., Wang, J., Yu, Y., Song, W., Yisimayi, A., Wang, J., An, R., Chen, X., Zhang, N., et al. (2022). Imprinted SARS-CoV-2 humoral immunity induces convergent Omicron RBD evolution. Nature. 10.1038/s41586-022-05644-7.

23. CDC COVID Data Tracker: Variant Proportions. (2023).

24. EG.5 Initial Risk Evaluation, 9 August 2023. (2023).

25. NextStrain. (2023).

26. Qu, P., Faraone, J.N., Evans, J.P., Zheng, Y.M., Yu, L., Ma, Q., Carlin, C., Lozanski, G., Saif, L.J., Oltz, E.M., et al. (2022). Durability of Booster mRNA Vaccine against SARS-CoV-2 BA.2.12.1, BA.4, and BA.5 Subvariants. N Engl J Med 387, 1329–1331. 10.1056/NEJMc2210546.

27. Qu, P., Evans, J.P., Faraone, J.N., Zheng, Y.M., Carlin, C., Anghelina, M., Stevens, P., Fernandez, S., Jones, D., Lozanski, G., et al. (2023). Enhanced neutralization resistance of SARS-CoV-2 Omicron subvariants BQ.1, BQ.1.1, BA.4.6, BF.7, and BA.2.75.2. Cell Host Microbe 31, 9–17.e13. 10.1016/j.chom.2022.11.012.

28. Zeng, C., Evans, J.P., Pearson, R., Qu, P., Zheng, Y.M., Robinson, R.T., Hall-Stoodley, L., Yount, J., Pannu, S., Mallampalli, R.K., et al. (2020). Neutralizing antibody against SARS-CoV-2 spike in COVID-19 patients, health care workers, and convalescent plasma donors. JCI Insight 5. 10.1172/jci.insight.143213.

29. San Filippo, S., Crovetto, B., Bucek, J., Nahass, R.G., Milano, M., and Brunetti, L. (2022). Comparative Efficacy of Early COVID-19 Monoclonal Antibody Therapies: A Retrospective Analysis. Open Forum Infect Dis 9, ofac080. 10.1093/ofid/ofac080.

30. Zeng, C., Evans, J.P., Qu, P., Faraone, J., Zheng, Y.M., Carlin, C., Bednash, J.S., Zhou, T., Lozanski, G., Mallampalli, R., et al. (2021). Neutralization and Stability of SARS-CoV-2 Omicron Variant. bioRxiv. 10.1101/2021.12.16.472934.

31. Qu, P., Faraone, J.N., Evans, J.P., Zou, X., Zheng, Y.-M., Carlin, C., Bednash, J.S., Lozanski, G., Mallampalli, R.K., Saif, L.J., et al. (2022). Differential Evasion of Delta and Omicron Immunity and Enhanced Fusogenicity of SARS-CoV-2 Omicron BA.4/5 and BA.2.12.1 Subvariants. bioRxiv, 2022.2005.2016.492158. 10.1101/2022.05.16.492158.

32. Smith, D.J., Lapedes, A.S., de Jong, J.C., Bestebroer, T.M., Rimmelzwaan, G.F., Osterhaus, A.D.M.E., and Fouchier, R.A.M. (2004). Mapping the Antigenic and Genetic Evolution of Influenza Virus. Science 305, 371–376. doi:10.1126/science.1097211.

33. Kaku, Y., Kosugi, Y., Uriu, K., Ito, J., Kuramochi, J., Sadamasu, K., Yoshimura, K., Asakura, H., Nagashima, M., Consortium, T.G.t.P.J., and Sato, K. (2023). Antiviral efficacy of the SARS-CoV-2 XBB breakthrough infection sera against Omicron subvariants including EG.5. bioRxiv, 2023.2008.2008.552415. 10.1101/2023.08.08.552415.

34. Wang, Q., Guo, Y., Zhang, R.M., Ho, J., Mohri, H., Valdez, R., Manthei, D.M., Gordon, A., Liu, L., and Ho, D.D. (2023). Antibody Neutralization of Emerging SARS-CoV-2: EG.5.1 and XBC.1.6. bioRxiv, 2023.2008.2021.553968. 10.1101/2023.08.21.553968.

35. Cao, Y.R. (2023). F456L-carrying XBB*, like EG.5, is rapidly rising. Meanwhile, XBB*+L455F+F456L is also growing fast. Some updates explaining their advantages: .

36. Wang, Q., Bowen, A., Ho, J., Zhang, R., Valdez, R., Stoneman, E., Gordon, A., Liu, L., and Ho, D.D. (2023). SARS-CoV-2 Neutralizing Antibodies Following a Second BA.5 Bivalent Booster. bioRxiv, 2023.2008.2013.553148. 10.1101/2023.08.13.553148.

37. Chen, C., Nadeau, S., Yared, M., Voinov, P., Xie, N., Roemer, C., and Stadler, T. (2021). CoV-Spectrum: Analysis of Globally Shared SARS-CoV-2 Data to Identify and Characterize New Variants. Bioinformatics 38, 1735–1737. 10.1093/bioinformatics/btab856.

38. Fujita, S., Uriu, K., Pan, L., Nao, N., Tabata, K., Kishimoto, M., Itakura, Y., Sawa, H., Kida, I., Tamura, T., et al. (2023). Impact of Imprinted Immunity Induced by mRNA Vaccination in an Experimental Animal Model. The Journal of Infectious Diseases. 10.1093/infdis/jiad230.

39. Pfizer and BioNTech Submit Applications to U.S. FDA for Omicron XBB.1.5-Adapted Monovalent COVID-19 Vaccine. (2023).

40. MODERNA FILES FOR FDA AUTHORIZATION OF ITS UPDATED COVID-19 VACCINE. (2023).

41. Will New COVID Vaccines Work Against EG.5? (2023).

42. Imai, M., Ito, M., Kiso, M., Yamayoshi, S., Uraki, R., Fukushi, S., Watanabe, S., Suzuki, T., Maeda, K., Sakai-Tagawa, Y., et al. (2023). Efficacy of Antiviral Agents against Omicron Subvariants BQ.1.1 and XBB. N Engl J Med 388, 89–91. 10.1056/NEJMc2214302.

