## Supplemental Table S1 and Figure S1, S2 for "Immune Evasion and Membrane Fusion of SARS-CoV-2 XBB Subvariants EG.5.1 and XBB.2.3"

| Table S1: Bivalent vaccinated HCW, BA.4/5-wave first responder, and XBB.1.5-wave first responder cohort information. | | | |
|---|---|---|---|
| | Bivalent Vaccinated HCWs (n=14) | BA.4/5-Wave First Responders/ Household Contacts (n=20) | XBB.1.5-Wave First Responders (n=8) |
| **Age in Years at Sample Collection [Median (Range)]** | 36 (25-48) | 44 (27-58) | 53 (38-64) |
| **Gender [n (% of Total)]** | | | |
| **Male** | 8 (57.1%) | 4 (20.0%) | 5 (62.5%) |
| **Female** | 6 (42.9%) | 15 (75.0%%) | 3 (37.5%) |
| **Unknown** | NA | 1 (5%) | NA |
| **Sample Collection Window** | Dec 2022-Early Jan 2023 | Mar 2022-Sept 2022 | Feb 2023-Early Aug 2023 |
| **Type of Vaccine [n (% of Total)]** | | | |
| **Unvaccinated** | NA | 17 (85.0%) | 4 (50%) |
| **3-dose Moderna** | NA | 2 (10.0%) | 1 (12.5%) |
| **3-dose Pfizer** | NA | 1 (5.0%) | 1 (12.5%) |
| **2-dose Pfizer + 1 dose Pfizer bivalent** | 1 (7.1%) | NA | NA |
| **3-dose Pfizer/Moderna + 1 dose Pfizer/Moderna bivalent** | 12 (85.8%) | NA | 1 (12.5%) |
| **4-dose Pfizer + 1 dose Pfizer bivalent** | 1 (7.1%) | NA | 1 (12.5%) |
| **Sample Collection Timing [Median (Range)]** | | | |
| **Days post 3rd dose for recipients of three doses** | NA | 158 (64-183) | 540 (513-567) |
| **Days post bivalent dose** | 66 (23-108) | NA | 220.5 (150-291) |
| **Infecting Variant** | | | |
| **BA.4** | NA | 4 (20%) | NA |
| **BA.5** | NA | 7 (35%) | NA |
| **Undetermined** | NA | 9 (45%) | NA |
| **XBB.1.5** | NA | NA | 8 (100%) |

Summaries of the demographic information for each of the cohorts used for neutralization experiments depicted in **Figure 2**. “NA” means the category is not applicable to the cohort.

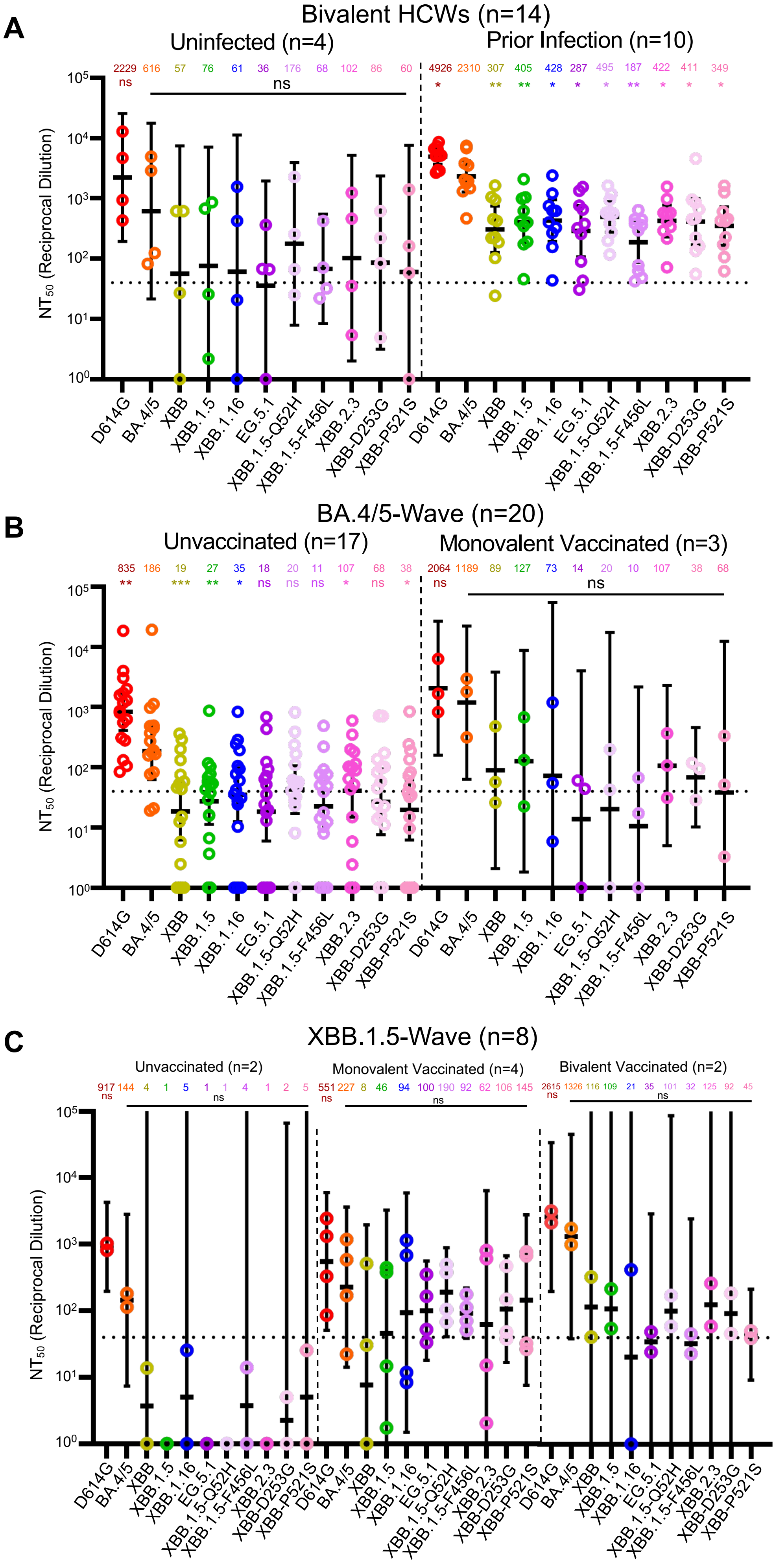

A
Bivalent HCWs (n=14)
Uninfected (n=4)
Prior Infection (n=10)
2229 616 57 76 61 36 176 68 102 86 60
ns
ns
4926 2310 307 405 428 287 495 187 422 411 349
* ** ** * * * ** * * *
NT50 (Reciprocal Dilution)
D614G BA.4/5 XBB XBB.1.5 XBB.1.16 EG.5.1 XBB.1.5-Q52H XBB.1.5-F456L XBB.2.3 XBB-D253G XBB-P521S
B
BA.4/5-Wave (n=20)
Unvaccinated (n=17)
Monovalent Vaccinated (n=3)
835 186 19 27 35 18 20 11 107 68 38
** *** ** * ns ns ns * ns *
2064 1189 89 127 73 14 20 10 107 38 68
ns
ns
NT50 (Reciprocal Dilution)
D614G BA.4/5 XBB XBB.1.5 XBB.1.16 EG.5.1 XBB.1.5-Q52H XBB.1.5-F456L XBB.2.3 XBB-D253G XBB-P521S
C
XBB.1.5-Wave (n=8)
Unvaccinated (n=2)
Monovalent Vaccinated (n=4)
Bivalent Vaccinated (n=2)
917 144 4 1 5 1 1 4 1 2 5
ns
ns
551 227 8 46 94 100 190 92 62 106 145
ns
ns
2615 1326 116 109 21 35 101 32 125 92 45
ns
ns
NT50 (Reciprocal Dilution)
D614G BA.4/5 XBB XBB.1.5 XBB.1.16 EG.5.1 XBB.1.5-Q52H XBB.1.5-F456L XBB.2.3 XBB-D253G XBB-P521S


**Figure S1**

**Figure S1: Subgroup analyses of infected and vaccinated individuals in neutralization data cohorts (Related to Figure 2).** Pseudotyped lentivirus bearing spikes of interest was used for a neutralization assay to determine neutralization titers for **(A)** bivalent mRNA vaccinated HCWs (n=14), **(B)** first responders and household contacts infected during the BA.4/5 wave in Columbus, OH (n=20), and **(C)** first responders infected during the XBB.1.5 wave in Columbus, OH (n=8). **(A)** The bivalent cohort was split into individuals who had no incidence of breakthrough infection before sample collection (n=4) and those that did experience breakthrough infection (n=10). **(B)** The BA.4/5-wave cohort was split into unvaccinated individuals (n=17) and individuals that received 3 doses of monovalent mRNA vaccine (n=3). **(C)** The cohort was divided into unvaccinated individuals (n=2), people that received 2 or 3 doses of monovalent vaccine (n=4), and people who received at least 3 doses of monovalent vaccine and a bivalent booster (n=2). Plots depict geometric mean neutralization tiers above each variant. The horizontal dashed line at NT50 = 40 represents the limit of detection for the assay. Significance was determined using a repeated measures one-way ANOVA with Bonferroni post-test within each group. Log10 transformed neutralization titers were used to determine significance throughout. p values are displayed as $*p < 0.05$, $**p < 0.01$, $***p < 0.001$, and ns $p > 0.05$.

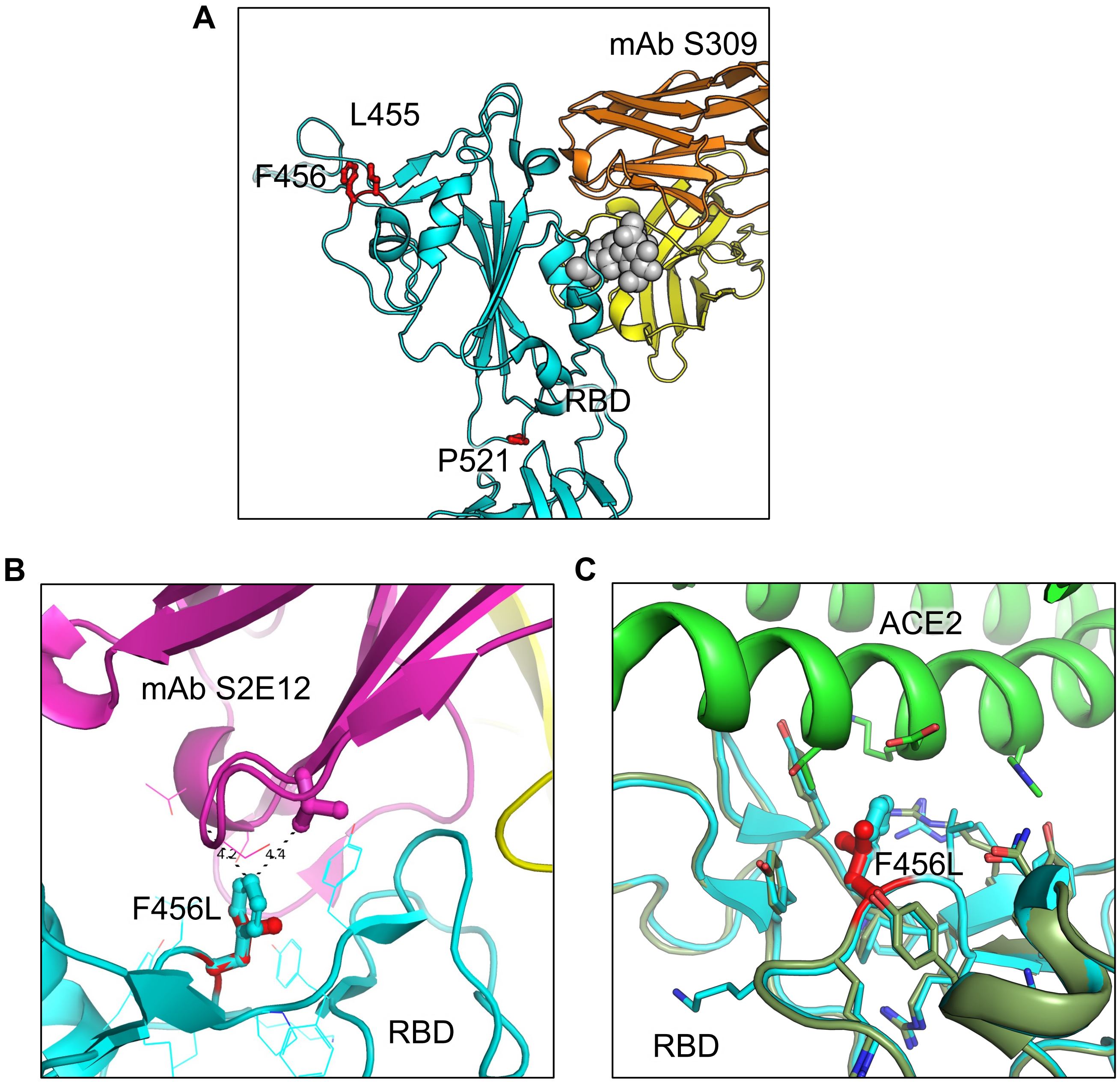
A
mAb S309
L455
F456
RBD
P521
B
mAb S2E12
4.2
4.4
F456L
RBD
C
ACE2
F456L
RBD

**Figure S2**

**Figure S2: Modeling of the F456L mutation at interface with ACE2 and mAbs S309 and S2E12.** Homology modeling was used to simulate the effects of the F456L mutation present in the EG.5.1 spike on **(A)** binding of class III monoclonal antibody S309, **(B)** binding of class I monoclonal antibody S2E12, and **(C)** ACE2 binding. **(A)** Spike RBD is depicted in cyan and mAb S309 is depicted in orange and yellow. Key spike mutations of EG.5.1 in positions F456 and P521 are highlighted in red. Glycan epitope for S309 on is depicted as grey space-filling spheres. **(B)** EG.5.1 spike is represented in cyan and mAb S2E12 in purple and yellow. Residue F456L is represented in the center with distances to residues Val52 and Gly54 represented by black dashed lines. **(C)** EG.5.1 spike is represented dark green overlaid D614G spike in cyan. The F456L mutation is highlighted in red. ACE2 is represented in green.
